# Estimation of the SNP mutation rate in two vegetatively propagating species of duckweed

**DOI:** 10.1101/2020.06.26.173039

**Authors:** George Sandler, Magdalena Bartkowska, Aneil F. Agrawal, Stephen I. Wright

## Abstract

Mutation rate estimates for vegetatively reproducing organisms are rare, despite their frequent occurrence across the tree of life. Here we report mutation rate estimates in two vegetatively reproducing duckweed species, *Lemna minor* and *Spirodela polyrhiza*. We use a modified approach to estimating mutation rates by taking into account the reduction in mutation detection power that occurs when new individuals are produced from multiple cell lineages. We estimate an extremely low per generation mutation rate in both species of duckweed and note that allelic coverage at *de novo* mutation sites is very skewed. We also find no substantial difference in mutation rate between mutation accumulation lines propagated under benign conditions and those grown under salt stress. Finally, we discuss the implications of interpreting mutation rate estimates in vegetatively propagating organisms.

## Introduction

Most research on the evolution of mutation rates has focused either on sexually reproducing eukaryotes or unicellular organisms, both of which feature a single cell phase as part of their life cycle. However, a diverse array of organisms reproduce either through clonal budding, fission or vegetative growth, whereby a single cell phase is not imposed every generation (Bell 1982). This mode of reproduction potentially allows multiple cell lineages to be transmitted from parent to offspring, complicating the process of genotyping individuals. This happens because when individuals composed of a mosaic of cells are sequenced, the mean number of sequencing reads supporting non-reference mutations is no longer 50%. Such a skew in allelic coverage makes it harder to distinguish true mutations from sequencing errors (Cibulskis *et al*. 2013), complicating the assessment of power when calculating per base pair mutation rates. Even if a cellular mutation rate can be calculated for an organism with multiple cell lineages, it becomes more challenging to use this parameter in population genetics analyses as mutations can potentially be lost within an organism before truly contributing to population level genetic diversity. Previous theoretical work modeling mutation load in organisms with multiple cell lineages has suggested that cell lineage selection can significantly reduce mutation load by purging deleterious mutations during somatic growth (Otto and Orive 1995). As most new mutations are thought to be deleterious (Sturtevant 1937; Eyre-Walker and Keightley 2007) this type of selection might skew the level of genetic diversity observed in organisms with vegetative reproduction compared to the level expected given their per base pair mutation rates.

Previous studies have investigated the rate of somatic mutations in plants where multiple cell lineages can segregate within a generation (Watson *et al*. 2016; Schmid-Siegert *et al*. 2017; Plomion *et al*. 2018; Wang *et al*. 2019). While somatic mutations can be transmitted from generation to generation in plants (Plomion *et al*. 2018; Wang *et al*. 2019), if somatic growth is followed by sexual reproduction, a single cell bottleneck is nonetheless imposed on any segregating variation within the soma, removing the persistence of multiple cell lineages across generations. This is however not the case for organisms reproducing through vegetative growth, budding or fission. Despite their frequency across the eukaryotic tree of life, almost no per-base-pair mutation rate estimates exist for organisms procreating through such modes of reproduction. One recent study in a vegetatively growing fairy-ring mushroom reported very low mutation rates per mitotic cell division (Hiltunen *et al*. 2019). The authors of this work used simulated mutations to assess the level of power they had to detect low frequency *de novo* mutation in this dataset, improving their estimate of the fairy ring mushroom mutation rate.

Here we report mutation rate estimates in two species of duckweed (*L. minor* and *S. polyrhiza*). Both species are free-floating, facultatively sexual aquatic plants. While duckweed can produce seed though sexual reproduction, most growth occurs vegetatively via clonal budding from two pouches present in the duckweed frond (Landolt, 1986). While these species are found all across the globe and likely have enormous census population sizes, allozyme and genomic analyses have revealed low levels of genetic diversity within local populations (Cole and Voskuil 1996; Ho 2018; Xu *et al*. 2019; Ho *et al*. 2019). Work by Xu *et al* (2019), has estimated the per base pair mutation rate in a genotype of *S. polyrhiza* grown in the field and the lab, finding an extremely low rate of mutation in both cases. However, their analysis did not take into account the fact that duckweed individuals are likely composed of a mosaic of cell lineages during periods of asexual growth, potentially leading to an underestimate of the true mutation rate.

Studying two duckweed species allows us to contribute to three other questions in mutation rate evolution research. First, our mutation rate estimates provide another species to add to the existing set of species with mutation rate estimates that, collectively, allow for testing the theory that selection against mutators should be most efficient in species with large effective population sizes (Sung *et al*. 2012; Lynch *et al*. 2016). Duckweeds are useful additions to this set as previous work has suggested that the effective population size (*N*_*e*_) of *S. polyrhiza* is on the order of a million individuals (Ho *et al*. 2019; Xu *et al*. 2019). Second, the inclusion of two facultatively sexual species that differ in their degree of sexuality allow us to preliminarily investigate the effect of recombination on the evolution of mutation rates. Genomic and allozyme patterns have suggested that *L. minor* undergoes bouts of sexual reproduction more often than *S. polyrhiza*, a pattern that is in line with flowering observations of these species in the field (Hicks, 1932; Landolt, 1986). Theoretical work has shown that when recombination breaks apart associations between mutator alleles (that elevate mutation rate) and the mutations they produce, mutation rates can evolve in several ways. On one hand selection against mutators in more sexual populations may be relaxed as they no longer remain linked to new deleterious mutations (Kimura 1967; Leigh 1970). Alternatively mutator alleles can spread when recombination is sufficiently low if they hitch hike along with any beneficial mutations they produce (André and Godelle 2006). Finally, environmental stress is known to increase mutation rates in bacteria in a process known as stress-induced mutagenesis (Foster 2007). A few examples of stress increasing mutation rates are known in eukaryotes (Matsuba et al. 2013; Jiang et al. 2014; Sharp and Agrawal 2016; but see Saxena et al. 2019), however it is unclear how general this phenomenon is. We performed our experiment both under a control and salt stress treatment to test whether stress-induced mutagenesis is a common phenomenon in plants.

We estimated the mutation rate in 46 asexually propagated mutation accumulation lines, including two genotypes of *S. polyrhiza* and one genotype of *L. minor*. We report an exceptionally low rate of mutation in both species of duckweed and note a pattern of skewed allelic counts at *de novo* sites that suggests the presence of multiple segregating cell lineages in vegetatively reproducing duckweed.

## Materials and Methods

### Mutation accumulation and DNA extraction

MA lines were started for three genotypes in April 2014 and propagated for approximately 60 generations. Two *Spirodela polyrhiza* genotypes were used: GP23 from Grenadier Pond, Toronto, Canada and CC from Cowan Creek, Oklahoma, USA. A single genotype of *L. minor* (GPL7) was also isolated from Grenadier Pond, Toronto, Canada. For each genotype (CC, GP23 and GPL7), we established 16 MA lines. We generated each line from a single maternal plant, which was started by isolating two fronds from each genotype culture. Because daughter fronds are generated iteratively, we grew and isolated daughters tracking pedigree until a minimum of 16 daughter fronds paired by generation were available for each of the three genotypes (arising from a single starting maternal frond). Daughters in each frond pairs (matched for generation relative to maternal frond) were assigned to one of two growth medium treatments (salt stress and control). Daughters are produced from two pockets of meristem tissue on either side of the maternal frond and mature daughter fronds remain attached to the maternal plant via a stipule for a short time (Landolt, 1986). To ensure that each generation was propagated with a daughter frond, we separated the daughter from the maternal frond as soon as the daughter began producing her own frond. Each line was checked for mature daughter fronds every two days. The first daughter produced was used whenever possible. MA lines were propagated in 0.5X Appenroth liquid growth medium (Appenroth *et al*. 1996) at 24°C with 12 hours of light per day. Generation times were similar in both species at ∼2.9 and ∼2.8 days under normal conditions and 3.5 and ∼3.3 days under salt stress for *S. polyrhiza* and *L. minor* respectively. Salt stress lines were supplemented with 25mM of NaCl for *S. polyrhiza* and 50mM of NaCl for *L. minor*. Prior to the start of the MA experiment, we performed growth assays to establish stressful NaCl levels for both species. The chosen salinity levels caused duckweed fronds to become patchy and thin but still allowed for continual asexual propagation.

After the termination of the MA experiment we allowed MA lines to continue growing for several generations without removing any individuals to obtain enough plant material to perform CTAB DNA extractions.

### Sequencing and filtering

We sequenced the MA lines at the McGill Innovation Centre. Illumina HISeq 2000 sequencing with 100bp paired end reads was used for both *S. polyrhiza* genotypes while Illumina HISeq 2500 sequencing with 125bp paired end reads was used for the *L. minor* genotype.

Paired end reads were mapped to the *S. polyrhiza* and *L. minor* reference genomes (Van Hoeck *et al*. 2015; Michael *et al*. 2017) using the Burrows-Wheeler aligner (BWA) 0.717 using the BWA-MEM option (Li and Durbin 2009). We then used Picard to remove duplicate reads before calling indels using the HaploTypeCaller tool in GATK 3.7 (McKenna *et al*. 2010). Next, we used the IndelRealigner tool in GATK to perform indel realignment. Finally, we used BCFtools (1.6) (Li 2011) to create mpileup files for the realigned output from GATK and to call SNPs and short indels (indels no more than 10bp). After mapping, mean and median coverage was 26, and 25 for individual *S. polyrhiza* lines, and 18, 17 for individual *L. minor* lines. We also calculated total median coverage for each site within each genotype (by summing across all individual lines), which was 436, 426, 280 for genotypes GP23, CC, and GPL7 respectively.

We first filtered out sites with unusually high or low coverage. We did this by eliminating sites that had coverage outside +/- 200x median coverage (summed across all lines) in each *S. polyrhiza* genotype and +/- 100x median coverage in the *L. minor* genotype due to the lower quality of the reference genome for this species. We visualised relatedness between our lines using a PCA plot created from heterozygous sites present in our MA lines in R (v5.3.5) (R Core Team 2019) using the package SNPRelate (Zheng *et al*. 2012). In doing so we discovered two major outliers in one of our *S. polyrhiza* genotypes (CC) suggesting that these two lines were cross contaminated. We subsequently removed these two lines from our analysis.

Our next round of filtering aimed to remove low quality regions of the genome that contain unusually high amounts of in-phase heterozygous variants (e.g. Figure S1). Such regions likely represent collapsed duplications in the reference genome that map poorly to an incorrect genomic coordinate. These variants are highly reference-biased in their allelic coverage likely due to the poor mapping of reads that contain many differences relative to the reference genome. To remove such regions, we first created a consensus genotype for each set of lines; if more than one line in a given genotype supported the existence of a heterozygote at a site, that site was designated as heterozygous in the consensus genotype. We then performed a sliding window analysis on heterozygosity on each consensus genotype. We used 1000bp windows with a 100bp step. After trying a variety of filtering criteria, we decided to designate regions of the genome as callable if there existed no more than 10 heterozygous calls in a 1000bp window in each *S. polyrhiza* genotype and no more than 5 heterozygous calls per 1000bp window in the *L. minor* genotype. We used more stringent criteria in *L. minor* due to the lower quality of the genome assembly. These cut-offs represent a trade-off between eliminating problematic, variant-rich areas of the genome and excluding well assembled genomic areas with higher than average diversity. After filtering, around 100Mb of the genome was retained as callable in each of the three genotypes. This filtering step greatly improved the allelic coverage of ancestral heterozygous sites by removing suspected hidden duplications that map poorly to the reference (Figure S2).

Next, we called putative *de novo* mutations in the remaining callable regions. Within each set of lines, we picked sites where one line had a heterozygous genotype, with at least 5 reads supporting the non-reference base, but all other lines supported a homozygous genotype. We then extracted such sites from the mpileup file used to call genotypes. This was done as the pileup file contains reads that are filtered out during genotype calling but are useful in our case as they can lead to the elimination of false positive mutations. We filtered putative *de novo* mutations using the mpileup file in two ways. First, if a line other than the one which contained the *de novo* mutation had any reads which supported the *de novo* base call we excluded the site. Second, if a site with a *de novo* base call contained reads with more than two non-reference bases across all samples, we also excluded the site. We did this to exclude sites where a high rate of sequencing errors might have occurred. We used this cut-off based on the observation that at sites where all lines supported a homozygous genotype, the vast majority of sites contain no more than one alternate base call (again likely due to sequencing errors which can be observed in the mpileup file). The remaining putative *de novo* mutations that passed these filtering criteria were visually inspected in the Integrative Genomics Viewer (IGV) (Robinson *et al*. 2011). We excluded a few mutations which appeared on reads in complete linkage with other non-reference bases (an indication of hidden genomic duplications) or on reads that looked like the product of PCR or sequencing errors (see Figure S3-5 for examples).

### Power analysis

To calculate the per generation mutation rate, we first needed to know how much power we had to detect *de novo* mutations at our callable sites. To assess power, we first obtained a list of sites where we knew we had non-zero power to detect mutations, this included sites where all lines within a genotype supported a homozygous reference base call and no more than one alternate base was present in the mpileup file (one less than in the case if a *de novo* mutation was present). We then randomly sampled 500,000 such sites from each genotype independently, and randomly chose a line where a mutation could have happened. We randomly eliminated a third of the sites where one alternate base was present in the mpileup file as our filtering criteria would eliminate true *de novo* mutations if another line by chance contained a sequencing error which matched the *de novo* base call (i.e., we assume the probability of this occurring is 1/3). Of the remaining sites, we assigned how many reads would support the *de novo* base call by drawing from a binomial distribution with a success rate of 50%, 34%, 28%, 20% and 10%. These different values were chosen to represent a range of frequencies a mutation may be found due to the inheritance of multiple cell lineages in asexual reproduction. This is similar to the approach taken by Hiltunen *et al*. (2019) when calculating mutation rates in vegetatively growing fairy-ring mushrooms.

Our estimate of power was the proportion of sites (out of the original 500,000) that had at least 5 reads supporting the *de novo* mutation (for each possible binomial success rate that we tested). We then multiplied our power estimates by the number of callable sites in the genome. Then separately for each line, we multiplied the adjusted number of callable sites by the number of MA generations and summed these values across all lines in a given genotype (split by treatment). This provided us with a denominator for our mutation rate calculation. Our final step was to divide the number of *de novo* mutations identified in each genotype (split by treatment) by this denominator. The *de novo* mutation count was adjusted for false positives identified during mutation validation.

### Calling indels

We scanned our lines for *de novo* indels in the same way as we searched for point mutations with one key modification. We used the same regions of the genome that we had previously assessed as “callable”; however, we only considered *de novo* indels if they were at least 2000 bp away from any other indel in any other line of the same genotype. This filtering step was required to avoid false positive indel calls which appear due to spurious mapping patterns in repetitive regions.

### Mutation validation

We only had tissue from *S. polyrhiza* to perform validation on putative *de novo* mutations as unlike *L. minor, S. polyrhiza* produces asexual resting stages called turions which we were able utilize for long term refrigerated storage. After allowing turions to germinate we extracted DNA from our MA lines using a Qiagen DNeasy kit. Afterwards, we designed primers for 14 SNPs and one indel found in our two *S. polyrhiza* genotypes. We performed PCR reactions using FroggaBio PCR mastermix which were then sent off for Sanger sequencing at Eurofins Genomics. We inspected the Sanger sequencing chromatograms in FinchTV 1.4.0 (Geospiza, Inc.; Seattle, WA, USA; http://www.geospiza.com) to see if we could detect a peak suggesting the presence of a mutant base in the focal MA line and checked that the base was not present in a different MA line of the same genotype as a negative control. One line from the *S. polyrhiza* CC genotype (line E) contained a massive overabundance of putative *de novo* mutations (of the 29 initial mutations found in 14 lines of the CC genotype, 9 were identified from this line); all 3 variants that we attempted to Sanger sequence validate from this line failed. Ultimately, we excluded this CC line from all the results presented here. Excluding CC line E, 11 total SNPs that were checked with Sanger sequencing and used to correct for the false positive rate in mutation detection in all our *S. polyrhiza* lines. We adjusted our mutation rate for false positive by excluding mutations that failed validation, summing all mutations that passed validation, and multiplying the sum of unvalidated mutations by our estimate of the false positive rate. Our formula for the true positive rate is as follows:

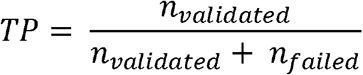

Where *n*_*validated*_ refers to the total number of mutations successfully validated in *S. polyrhiza*, and *n*_*failed*_ refers to the total number of mutations that failed validation.

Finally, the formula for our per generation, per base pair mutation rate estimates is as follows:

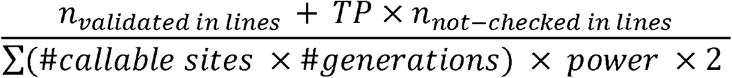

where *n*_*validated in lines*_ refers to the number of validated mutations in the focal set of lines, and *n*_*not-checked in lines*_ refers to the number of mutations not tested with Sanger sequencing in the focal set of lines. The number of mutations and callable sites was summed across all lines within a genotype, within a treatment. The true positive rate, TP, was estimated once using all Sanger-tested mutations from across the entire experiment.

### Statistical analysis

We calculated 95% confidence intervals for our mutation rates using the Agresti Coull method implemented in the R package “binom” (Dorai-Raj 2014). We tested for significant differences between our salt and normal treatment lines using independent Chi-square tests in each genotype. To test for differences between *L. minor* and *S. polyrhiza* we merged the number of callable sites (scaled by power) and number of mutations in the two *S. polyrhiza* genotypes (separately for each treatment), and also applied C hi-square tests to test for significance in mutation rate variation between species and conditions.

### Data availability

Raw sequencing data is available on the NCBI SRA database under accession PRJNA659313 for *S. polyrhiza* and PRJNA659264 for *L. minor*. Custom scripts and additional data are available at https://github.com/gsan211/duckweed_MA. Supplementary material has been uploaded as a separate file.

## Results

### De novo mutations

In total, we identified 23 and 19 putative *de novo* mutations in *S. polyrhiza* genotypes GP23 and CC, respectively and a further 29 mutations in *L. minor* (Supplemental table 1). When inspecting the putative *de novo* mutations in IGV, we observed that most mutant sites exhibited highly reference biased allelic counts (over 50% of reads support the reference base call). On average, mutant sites contained 26%, 30%, and 34% mutant reads with SD 8.2%, 11.6%, and 9.1% in genotypes GP23, CC (*S. polyrhiza*) and GPL7 (*L. minor*) respectively. We implemented several filtering and quality control steps to ensure these mapping patterns were not a result of sequencing error or genome mis-assembly. First, we masked areas of the genome that had odd coverage patterns or were enriched for heterozygous calls to avoid areas containing hidden genomic duplications. The heterozygous variants that pass these filtering criteria and are present in the ancestral genotypes (i.e. are shared by all MA lines within a genotype) show relatively normal patterns of reference and non-reference allele coverage. Second, we only considered *de novo* mutations at sites with minimal sequencing errors. This step eliminated problematic areas of the genome prone to genotyping error. Finally, we performed Sanger sequencing validation on 11 putative *de novo* mutations (in *S. polyrhiza*) of which 6 were validated with both positive and negative controls. The pattern of reference bias in our *de novo* mutations was also present in our Sanger sequencing chromatograms; the reference base peak was generally much larger than the mutant peak. When we plotted Illumina mutant base frequency vs. Sanger mutant peak height (standardized by reference peak height), we observed strong concordance for our 6 validated mutations (R^2^ = 0.56; Figure 1) suggesting that these are not spurious mapping patterns or sequencing errors but rather reflect a true bias in mutation abundance. When we inspected our PCR products with gel electrophoresis we observed single, clean bands of the expected product size so we consider it unlikely that multiple sites in the genome were amplified leading to odd mapping patterns at our *de novo* mutation sites. One alternative is that reference biased mapping patterns may be due to some bias in the PCR amplification process. We do not believe this is likely however since our primers were designed for sequences that were flanking the site of *de novo* mutations that should be identical whether a *de novo* mutations is present or not.

**Figure 1.**
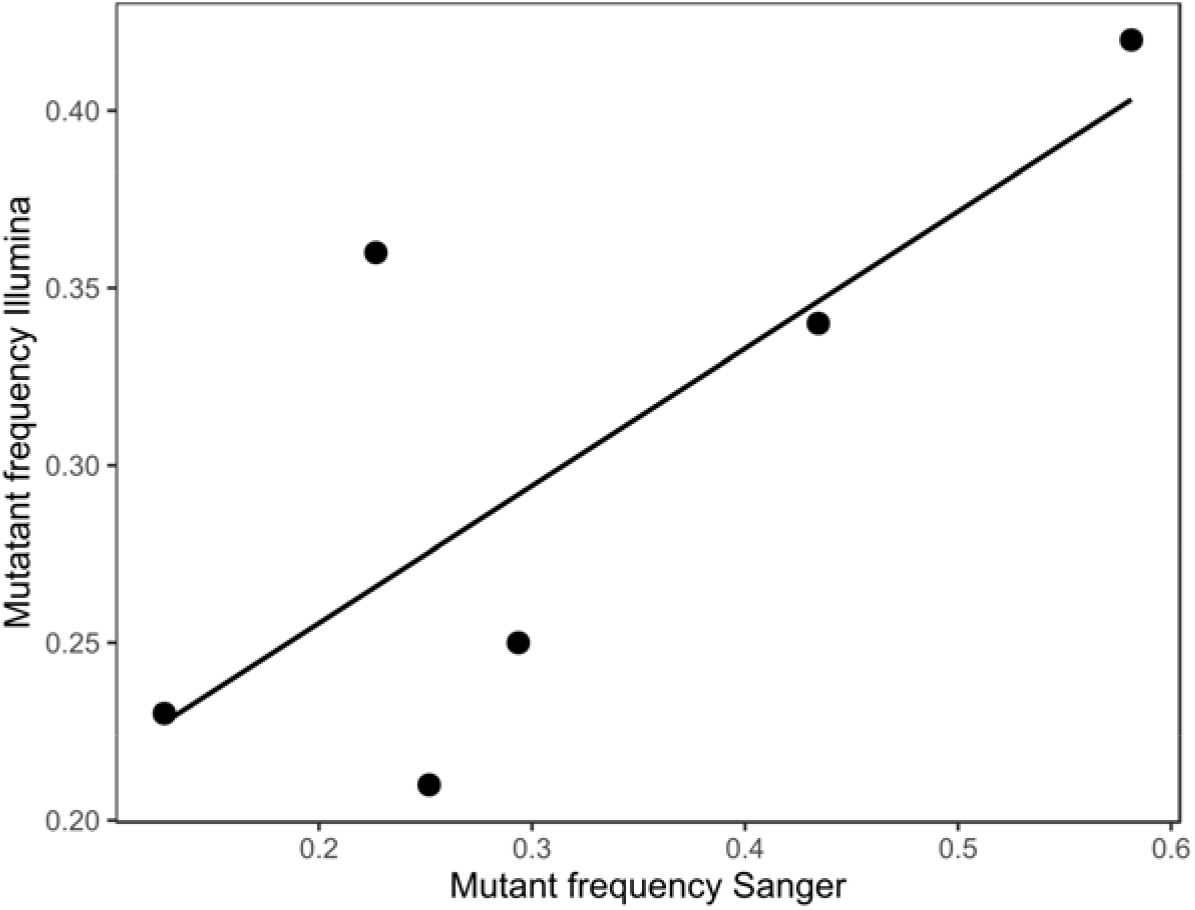
Proportion of de novo bases from Illumina reads vs. relative peak height of mutant base in Sanger sequencing data in Spirodela polyrhiza. R^2^ = 0.57.

We report a high false positive rate for *de novo* mutation identification in *S. polyrhiza* of ∼45%. This is likely a consequence of having to distinguish between true mutations with low allelic counts and sequencing errors or bioinformatic artifacts that can appear at similarly low frequencies. Additionally, background noise in Sanger Sequencing chromatograms can obscure variants with low allelic counts, posing a potential way in which false positive rates may be artificially elevated. We attempted to avoid this issue by ensuring that our chromatograms had relatively low levels of background noise before confidently assigning mutations as failing or passing validation.

The majority of mutations were C -> T transitions in both species of duckweeds (Figure 2) in concordance with patterns found in previous mutation rate studies (Ossowski *et al*. 2010; Thomas *et al*. 2018). Transitions were more common than transversions, with the average ti/tv ratio being 2.23 for *S. polyrhiza* and 4.5 for *L. minor*. This is consistent with previous results reported in *S. polyrhiza* by Xu *et al*. (2019) where three C -> T and one C -> A mutations were detected and validated. Approximately 50% of C -> T transitions occurred at CpG sites in both species, a pattern that is consistent with previous evidence of elevated mutation rates at such sites (Hodgkinson and Eyre-Walker 2011). We used SNPeff (Cingolani *et al*. 2012) to annotate our putative *de novo* mutations with most SNPs being labelled as intergenic (See supplemental table 1).

**Figure 2.**
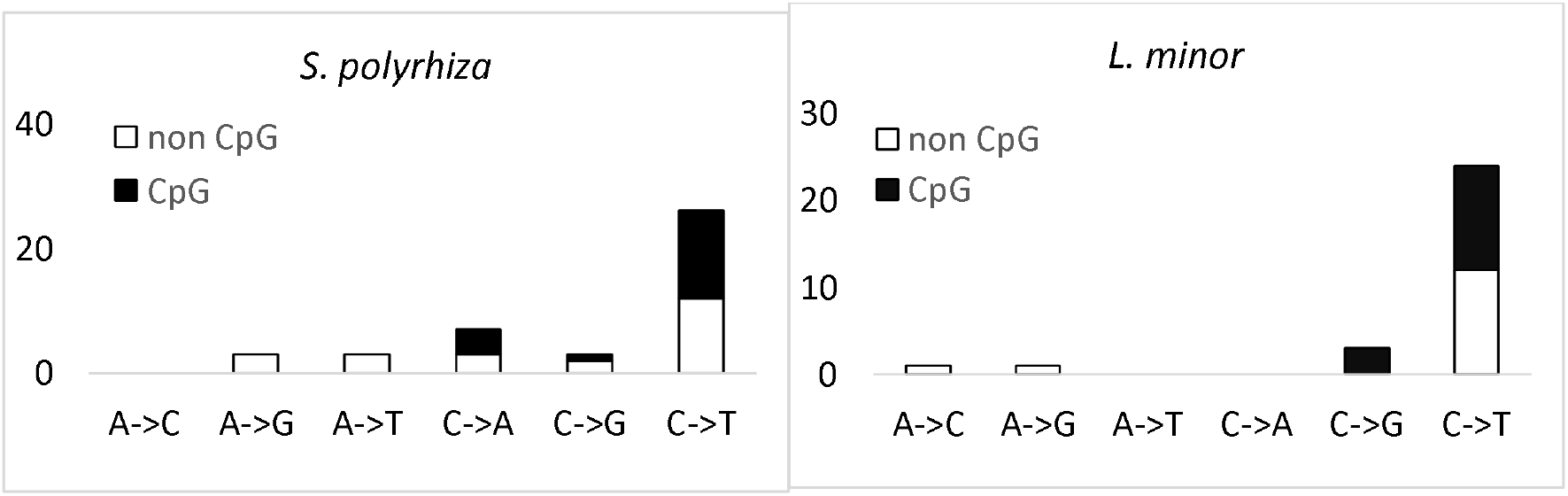
De novo mutation spectra in two species of duckweed (Spirodela polyrhiza, Lemna minor).

Our analysis uncovered only one putatively *de novo* short indel that turned out to be a false positive based on our Sanger sequencing validation analysis.

### Mutation rate comparisons between *S. polyrhiza* and *L. minor*

Our estimate of the per generation, per base pair SNP mutation rate highly depends on our power to detect mutations, which in turn depends on factors such as total read depth and the number of reads that support a *de novo* mutation base-call (See Supplemental tables 2-5). Heterozygous SNPs are expected to be supported by around 50% of reads in organisms with a single cell phase. In our case, on average, 28% (*S. polyrhiza*) and 34% (*L. minor*) of reads supported *de novo* heterozygous mutations. To improve our estimate of the mutation rate, we assumed that we should only expect to find mutations that are on average supported by 28% (in *S. polyrhiza*) or 34% (in *L. minor*) of reads at each mutant site (see methods for more details of power analysis). In this case, our point estimate of the mutation rate for plants grown in standard medium is 8.39E-11 for *S. polyrhiza* and 8.66E-11 for *L. minor* (Figure 3). For plants grown in stressful salt medium our point estimates for the mutation rate are 9.91E-11 for *S. polyrhiza* and 16.9E-11 for *L. minor*.

**Figure 3.**
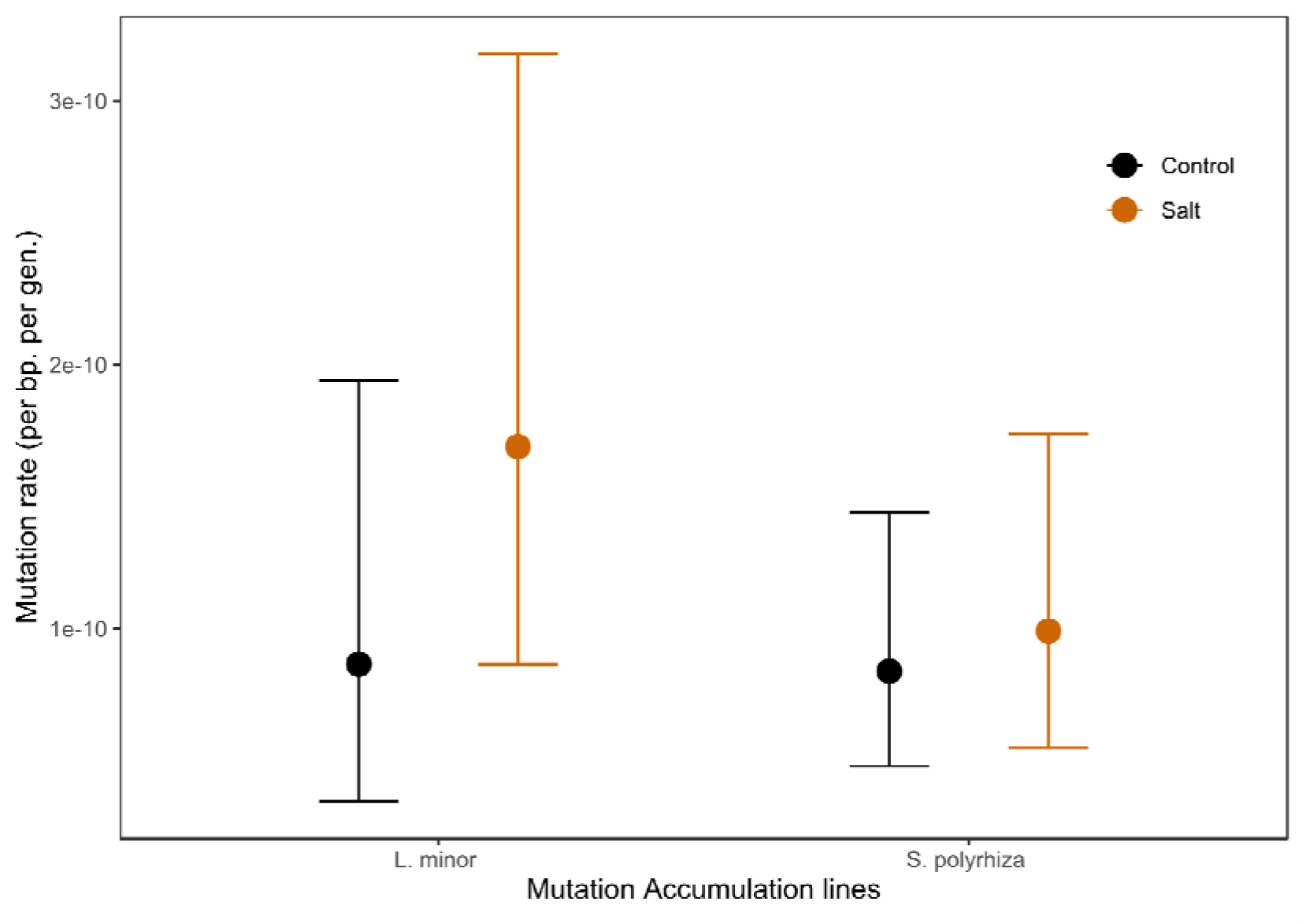
Mutation rate estimates in two species of duckweeds duckweed (Spirodela polyrhiza, Lemna minor). Error bars show Agresti-Coull 95% confidence intervals.

While the estimated mutation rate is higher for *L. minor* in both conditions, the difference between the two species is non-significant (chi-square test: p = 1 and p = 0.30 for control and salt stressed conditions). The difference in mutation rate in the two species is, however, very sensitive to the expected frequency of reads that support the mutant base. For example, if our power estimate is based on the assumption that *de novo* mutations are expected to constitute only 10% of reads, the mutation rate under normal conditions increases to 5.13E-10 in *S. polyrhiza* and 16.8E-10 in *L. minor*. Our mutation rate estimates were not significantly different between our two individual *S. polyrhiza* genotypes (chi-square test: p = 0.50 and p = 0.77 for control and salt stressed conditions). Supplemental tables 3-4 give the inferred mutation rates with alternative assumptions for the expected proportion of non-reference reads at sites harbouring *de novo mutations*.

### Effect of salt stress on mutation rate

There were no significant differences in the mutation rate between lines propagated in normal and salt stress medium for either of our three genotypes (Figure 4, chi-square test: p = 1.00, p = 0.53, p = 0.29 for genotypes GP23, CC, GPL7 respectively). There was also no consistent trend in the change of mutation rate with the addition of salt stress as genotype GP23 appeared to have a decreased mutation rate while genotypes CC and GPL7 appeared to have an increase in the mutation rate with the addition of salt stress.

**Figure 4.**
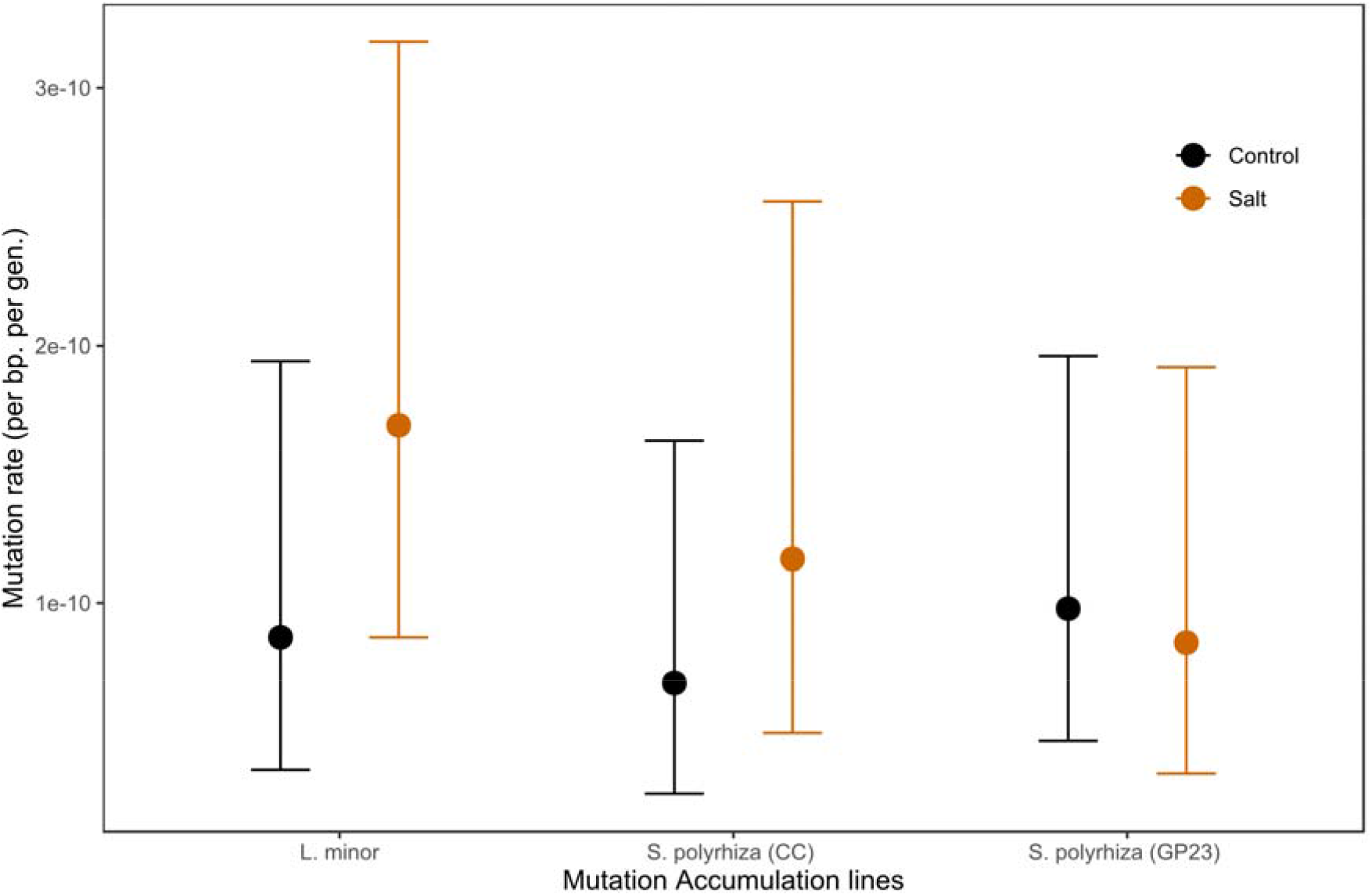
Effect of salt stress on the mutation rate in two genotypes of Spirodela polyrhiza and one genotype of Lemna minor. Error bars show Agresti Coull 95% confidence intervals.

Salt stress also did not have a significant impact on the ti/tv ratio. In two of the genotypes (1.33 -> 2 *S. polyrhiza* CC and 4.5 -> 8 *L. minor*), salt stress increased the ti/tv ration. In *S. polyrhiza* genotypes GP23 salt stress decreased the ti/tv ration (3.67 -> 2.00) although the difference was not significant in either of the three genotypes (chi-square test: p > 0.5 all cases). These results run counter to previous work in A. *thaliana* where salt stress was found to increase the de-novo mutation rate ∼two-fold and lower the ti/tv ratio (Jiang *et al*. 2014).

## Discussion

Using whole genome sequencing on 46 MA lines, we report a low per base pair, per generation SNP mutation rate in two species of duckweeds under two growth conditions. An important result in our study is that *de novo* mutations appear to have considerably reference biased genomic coverage in both duckweed species. We believe that this pattern is not indicative of sequencing or genome assembly errors but rather is a by-product of vegetative reproduction for several reasons. First, upon our inspection of the data in an independent study of mutation rates in *S. polyrhiza* by Xu et al.; we noticed similar levels of reference bias in Illumina short-read sequencing data at validated *de novo* mutations. Second, our own validation with Sanger sequencing showed that mutations with a higher reference bias in the Illumina dataset tended to have higher reference base peaks in their Sanger sequencing chromatograms and vice versa. Moreover, after filtering, most ancestral heterozygous sites that were shared by all lines in the three clonal genotypes were not reference biased in such a manner and those that were, were found in regions highly enriched for non-reference base calls. None of our putative *de novo* mutations are found in such regions; due to the low genetic diversity in duckweed, *de novo* mutations were generally the only heterozygous variants present in the immediate genomic area. This means that mapping bias due to divergence from the reference is unlikely to cause the strong allelic coverage bias we observed. Finally, a study of mutations in a vegetatively growing fairy-ring mushroom also reported patterns of reference bias with an average allelic coverage of 44% across 111 *de novo* mutations (Hiltunen *et al*. 2019). Therefore, the reference bias in our mutations is most likely a signature of the segregation of multiple cell lines from generation to generation in our duckweed MA experiment. Given that vegetative reproduction through clonal budding is the main form of reproduction in duckweed (Landolt, 1986), it seems reasonable that duckweed might not undergo a single cell phase for prolonged periods of time.

Simple models can be helpful in clarifying how multi-cell descent during vegetative reproduction may affect mutation rates and our estimation of them. Assume that clonal reproduction occurs by *n* parental cells forming a diploid offspring (generation 1). If a mutation occurs in one of these *n* cells, because of multi-cell descent, the offspring will be a genetic mosaic (i.e., not all cells will be genetically identical). The mutation begins at a frequency of 1/2*n* in the offspring. With conventional single-cell descent, where *n* = 1, a new mutation is expected to be at 50% frequency, but the frequency will be lower than 50% whenever *n* > 1. The n cells multiply in some (unknown) growth pattern to produce the mature offspring, which then itself reproduces. In the absence of detailed knowledge of developmental growth trajectories, we do not know what the representation of the original n cell lineages will be in the subsequent generation.

We can consider two simple scenarios: If only one of these original cell lineages gives rise to the next set of n cells used to produce the next generation (generation 2), then the mutation will either be completely lost from the lineage (if it was not in the chosen cell lineage) or it will be present in heterozygous state in all cells of future generations (if it was present in the chosen cell lineage). In this first scenario, the genetic mosaicism resulting from multi-cell descent persists only a single generation. Thus, in an experiment such as ours, only mutations occurring in the very last generation will be affected by this issue (i.e., multi-cell descent is a trivial issue in this case). A second scenario is that all cell lineages grow at equal rates and, each generation, *n* cells are chosen at random to form the next daughter. This is conceptually similar to the process of coalescence in a population (of cells here rather than individuals) of size *n*. Thinking backwards in time, all cells must eventually trace their coalescent history to a single cellular ancestor, but it may take many generations for this to happen (i.e., 2*n* generations, on average, but with variance proportional to *n*^2^). Thinking forwards in time, a mutation occurring in one of the n cells forming the generation 1 offspring may be present for many future generations, possibly becoming quite common within individuals, before eventually being present in all future cells (with chance 1/*n*) or none of them (with chance (*n* - 1)/*n*). With even modest values of *n* (e.g., 8 < *n* < 80), genetic mosaicism can persist for many generations. Mutations that will eventually be present in all lineages—as well as those that will eventually be eliminated—will thus be found below the 50% frequency expected in a “traditional” (i.e., non-mosaic) heterozygote. Although this second scenario as formulated here is unrealistically simple (e.g., random and independent choice of *n* cells for each generation), it illustrates how multi-cell descent can have consequences across multiple generations. Though developmental growth trajectories in duckweed are insufficient to formulate a more realistic model, we suspect that multiple cell lineages persist across multiple generations in duckweed and is responsible for the clear bias towards mutant SNPs being less than 50%. Recent work on segregating mutations in a single Zostera marina seagrass clone has made similar arguments to this model (Yu et al., 2020). In their study, Yu *et al*. uncovered a large class of reference biased SNPs that were present in some but not all sampled *Z. marina* ramets. By reconstructing the genealogical relationship of the sampled ramets, the authors demonstrated that such SNPs changed in frequency during vegetative growth, with some reaching heterozygous fixation in specific ramets. The authors argued that this data was well explained by a model of “somatic drift” whereby *de novo* mutations arise at low frequency within a single cell lineage before ultimately either reaching fixation or being lost.

Calculating mutation rates in organisms with multiple segregating cell lineages poses a technical challenge due to the difficulty of assessing power when mutations are present in only a subset of the cells of an individual. We implemented a method that takes into account the fraction of reads we expect to support a *de novo* mutation by using observed mapping patterns of putatively *de novo* mutations, similar to the approach used by Hiltunen *et al*. (2019). This method considers the fact that we have reduced power to detect recently arisen mutations that are at low frequency within their MA lines, giving a more accurate estimate of the SNP mutation rate. This approach is an admittedly crude attempt to address the problem. Rather than assuming the expected frequency of the mutant allele is 50%, we simply choose a lower value based on the observed coverage at putative *de novo* mutations. In reality, this lower value is unknown and will differ among mutations depending upon when each mutation arose, i.e., there is an unknown distribution of mutation frequencies at the end of the MA experiment. Nonetheless, our approach is an improvement over completely ignoring the issue. Moreover, the variation in mutation rate estimates inferred using different values for the assumed expected mutant frequency provides some sense of the sensitivity of these estimates to this assumption. However, the issue of the unknown distribution of mutation frequencies adds a caveat to the between-species comparison. Because of the moderate difference in coverage between species, which affects the power to detect mutations segregating at different true frequencies, the estimates for the two species may be somewhat differentially affected by any bias introduced by our method.

A natural consequence of the lack of a single cell phase is that from a population genetics perspective, the mutation rate becomes a harder parameter to interpret. On one hand, we may be interested in calculating the mutation rate that captures every new mutation that has arisen in a clonal bud. Alternatively, from an evolutionary perspective, we might be only be interested in mutations that will not only arise in a clonal bud but also persist in a future clonal descendant such that they contribute to population level genetic diversity. Aside from random inter-cell lineage “drift”, cell lineage selection may bias which *de novo* mutations are eventually lost in a clonal lineage (Otto and Orive 1995), although this process likely has a minimal effect on mutation rate estimates in most mutation accumulation studies. In principle, we might be able to calculate an “evolutionarily relevant” mutation rate by performing long term mutation accumulation experiments allowing the majority of *de novo* mutations to either be lost or to have fixed within a clonal lineage such that all cells in any clone will be either fully homozygous (in the case of loss) or heterozygous (in the case of fixation). However, from a practical standpoint, it is hard to know a priori how many generations will be necessary for this process to occur. In practice, we could attempt to estimate the frequency of *de novo* mutations that have become sufficiently common such that they are likely to not be lost before fixation leading to an estimate of an evolutionarily relevant mutation rate. For example, we could assume that mutations found in at least 50% of cells (i.e. at a frequency of at least 25% in a clonal bud) are more likely than not to eventually fix in their clonal lineage, being found in 100% of cells in some future generation. In reality, some of these mutations are still likely to be lost before they have a chance to fix while simultaneously some mutations that are below 50% frequency might still reach fixation in the future. More generally, the lower the frequency cut-off we use for mutations that are “likely” to fix, the better the chance that we capture all of the mutations that will reach fixation with the caveat that we will also be capturing more mutations that will eventually be lost. In our study we opted to use a read number cut-off rather than a frequency cut-off as we were primarily concerned about differentiating true *de novo* mutations from false positives, a task that is particularly challenging in organisms with low mutation rates. In practice, our filtering criteria results in us mostly identifying mutations with an allelic frequency of at least 20% (61/71 *de novo* mutation reported in our study). As mentioned above however, some of these mutations may still be lost prior to fixation within an organism meaning that the mutation rate we report here is likely inflated compared to a true “evolutionarily relevant” rate. To some extent, this upward bias is counter-balanced by mutations that could eventually go to fixation but are at too low frequency at the time of sequencing to be either observed or even inferred by our power calculation that is based on some threshold frequency (10-50%; Tables S2-5).

Our estimate of the mutation rate in *S. polyrhiza* (8.39E-11 per bp per gen.) is similar to the estimates reported by Xu *et al*. (7.92E-11 per bp per gen.), however there are two important differences between our studies. First, our MA experiment was conducted only in the lab, while Xu et al. placed MA lines both in the lab and in the field, observing no mutations in the lab setting, likely due to the smaller number of MA generations in their study. Second, Xu et al. used ancestral heterozygous sites to estimate their power to detect *de novo* mutations which are not strongly reference biased in a similar manner. This suggests that the estimate from Xu et al., while conducted in a more naturally realistic environment, may be an underestimate of the mutation rate in the field.

The estimates of the per generation, per base pair mutation rate in *S. polyrhiza* and *L. minor* are among the lowest so far for multicellular eukaryotes (Figure 5 and Table S6). One potential explanation for why duckweeds have lower mutation rates than other plants is the smaller number of cell divisions they likely go through compared to their larger, long-lived relatives. Mutation rate studies in trees have indeed shown that while per generation tree mutation rates are high (on the order of 1E-08 mutations per bp), mutation rates per unit growth (a proxy for number of cell divisions), must be several orders of magnitude lower (Xie *et al*. 2016; Hanlon *et al*. 2019; Orr *et al*. 2020). However, duckweeds do appear to exhibit a low mutation rate compared to animals which have limited cell divisions between meiotic events due to a segregated germline. Moreover, our per generation duckweed mutation rate estimates fall on the lower end of values seen in green algae and other unicellular eukaryotes, organisms which unlike duckweed only go through a single cell division per generation (Figure 5). Overall, these patterns fit with previous work that has suggested that number of cell divisions per generation alone is not enough to fully explain variation in mutation rates (Lynch 2010).

**Figure 5.**
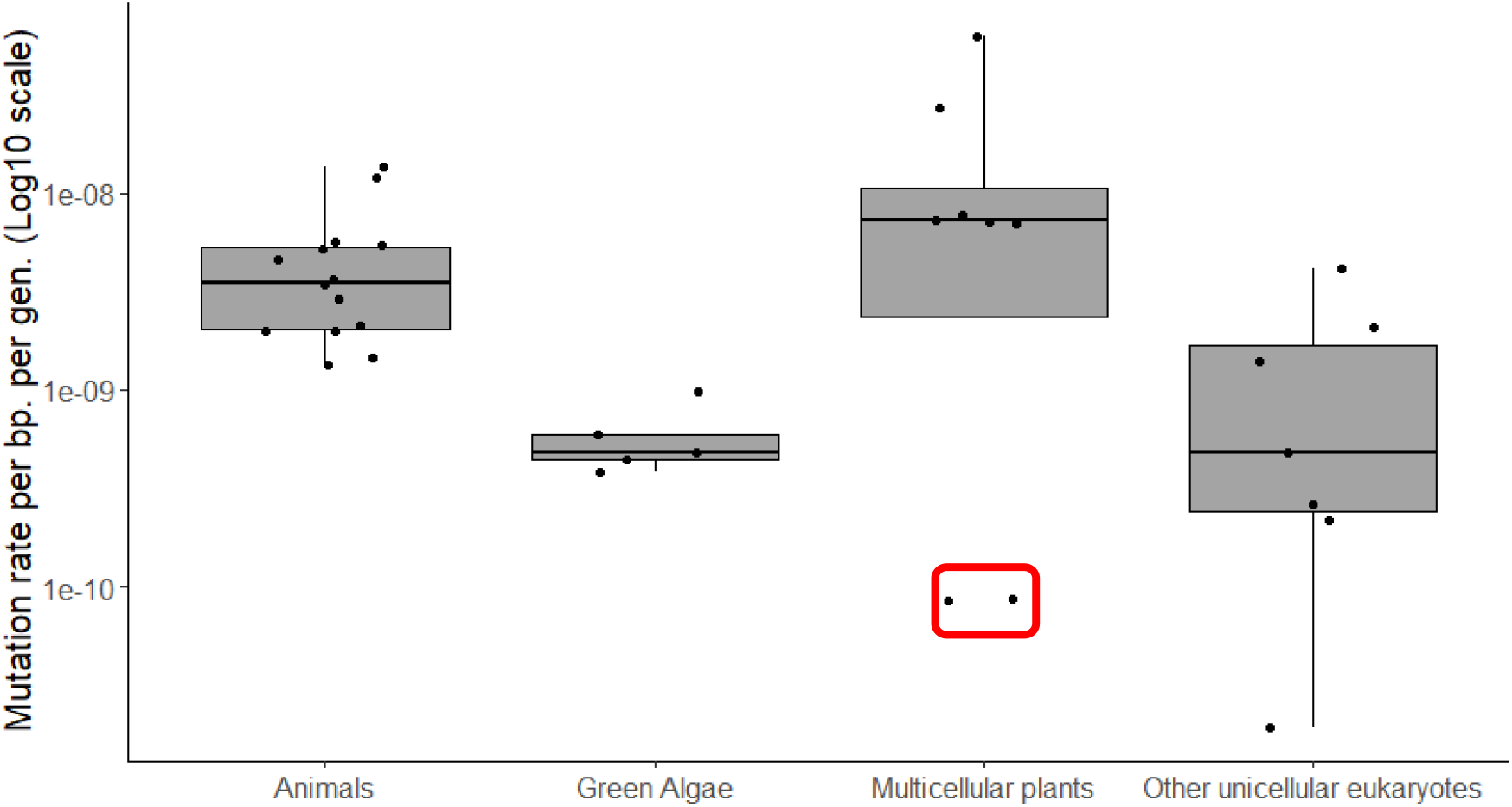
*Comparison of per base pair, per generation mutation rates between duckweed and other eukaryotic species. Duckweed estimated highlighted by red box. References*: (Ossowski *et al*. 2010; Lynch 2010; Denver *et al*. 2012; Schrider *et al*. 2013; Weller *et al*. 2014; Zhu *et al*. 2014; Venn *et al*. 2014; Keightley *et al*. 2015; Uchimura *et al*. 2015; Yang *et al*. 2015; Farlow *et al*. 2015; Ness *et al*. 2015; Smeds *et al*. 2016; Xie *et al*. 2016; Besenbacher *et al*. 2016 p.; Feng *et al*. 2017; Liu *et al*. 2017; Oppold and Pfenninger 2017; Flynn *et al*. 2017; Krasovec *et al*. 2017, 2018, 2019; Hanlon *et al*. 2019; Orr *et al*. 2020)

The low mutation rate observed in our study is consistent with efficient selection against mutator alleles in highly asexual organisms such as duckweed, bacteria and unicellular eukaryotes (Kimura 1967; Leigh 1970). This explanation would also predict that the mutation rate should be higher in *L. minor* than in *S. polyrhiza* as genomic analyses and field observations suggest that *L. minor* outcrosses more frequently than *S. polyrhiza* (Vasseur *et al*. 1993; Ho 2018; Xu *et al*. 2019; Ho *et al*. 2019). However, we did not observe a significant difference in mutation rate between species in our study. This could either be because we did not have the power to differentiate between such overall low mutation rates, because the difference in rates of sexual reproduction is not large enough between duckweed species, because a difference in rates of sex has arisen in recent history, or because factors other than reproductive mode play a larger role in shaping mutation rate evolution. The overall low mutation rate in both species is in contrast to the theoretical prediction that strong linkage in asexual genomes can allow mutators to fix in asexual populations if they hitchhike to fixation with beneficial mutations they produce (André and Godelle 2006). Population genomic analyses in duckweed have shown that selection on protein coding genes is weak as evidenced by elevated measures of π_N_/π_S_ (Ho 2018; Xu *et al*. 2019; Ho *et al*. 2019). This suggests that strongly beneficial mutations might be too rare to allow mutators to be selected for in duckweed.

Xu et al. inferred a global *N*_*e*_ for *S. polyrhiza* of ∼1×10E-06 so it might also be possible that the large effective population sizes of these species allow selection to be efficient enough to lower the mutation rate more than in most multicellular eukaryotes (Sung *et al*. 2012; Lynch *et al*. 2016). This explanation, however, is also inconsistent with the fact that measures for the efficacy of selection suggest that selection is weak in duckweed, likely due to the predominance of asexual reproduction (Xu *et al*. 2019; Ho *et al*. 2019).

In conclusion, we report a very low SNP mutation rate in two species of duckweed consistent with previous results in this group. We found that *de novo* mutations appear at low frequencies within MA lines suggesting the presence of multiple segregating cell lineages. We then used an approach that allows us to estimate the mutation rate when multiple cell lineages are transmitted across generations. The low mutation rate of these duckweeds is consistent with the idea that a higher degree of asexual reproduction leads to strong selection for low mutation rates.

## Acknowledgements

This work was supported by the Natural Sciences and Engineering Research Council of Canada (AFA and SIW). We thank Mitch Cruzan for helpful suggestions with data analysis. We thank Jade Lavallee and Victor Mollov for help with maintaining MA lines.

## Author contributions

SIW and AFA conceived the project. AFA, SIW, and MB collected samples. MB performed the MA experiment. GS did bioinformatics analyses. AFA, SIW, GS wrote the manuscript. All authors contributed to manuscript editing. SIW and AFA contributed equally to this work.

## Supporting Information

**Figure S1** Visualization of a low-quality genomic region that was filtered from our analyses.

**Figure S2** Allelic coverage at heterozygous sites before and after filtering.

**Figure S3** Visualization of a validated *de novo* mutation from both Illumina short read and Sanger sequencing data.

**Figure S4** Visualisation of a putative *de novo* mutation that was identified as a likely sequencing error

**Figure S5** Visualisation of a putative *de novo* mutation that was identified as a likely genome misassembly error

**Table S1** List of *de novo* mutations identified in two species of duckweed

**Table S2** Mutation rate estimates under the observed allelic bias at *de novo* mutations

**Table S3** Mutation rate estimates under several scenarios of allelic bias at *de novo* mutant sites in salt stressed mutation accumulation lines

**Table S4** Mutation rate estimates under several scenarios of allelic bias at *de novo* mutant sites in control mutation accumulation lines

**Table S5** Number and fraction of callable sites under several scenarios of allelic bias at *de novo* mutant sites

**Table S6** Summary of mutation rate estimates retrieved from the literature

## Notes

### Competing Interest Statement

The authors have declared no competing interest.

