## Supplemental figures and tables for "Estimation of the SNP mutation rate in two vegetatively propagating species of duckweed"

**Supplementary Tables and Figures**

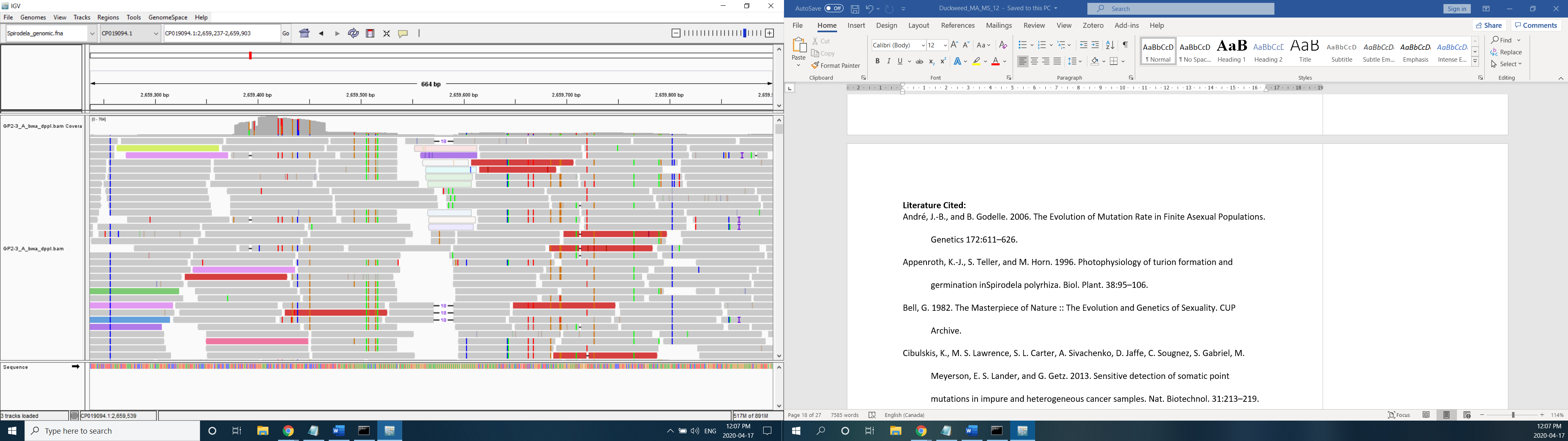

**Figure S1** Example of a region that was filtered out as a suspected hidden genomic duplication in the *S. polyrhiza* genome. Note the clusters of derived variants all present on the same reads.

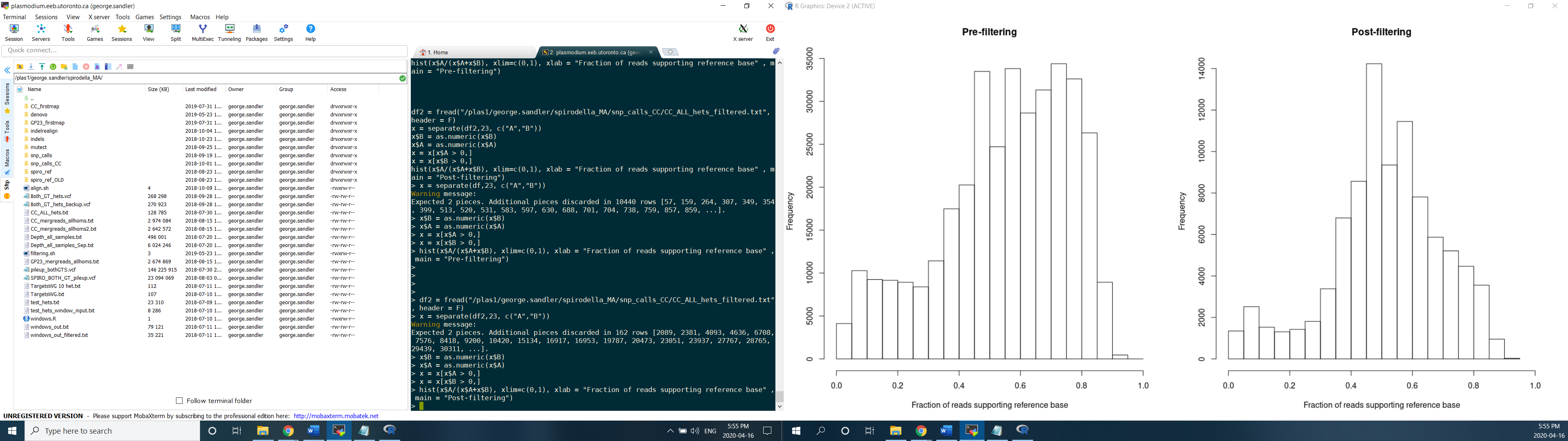

**Figure S2** Fraction of reads supporting reference base at ancestral heterozygous sites in line CC-F before and after genomic filtering of poor-quality regions.

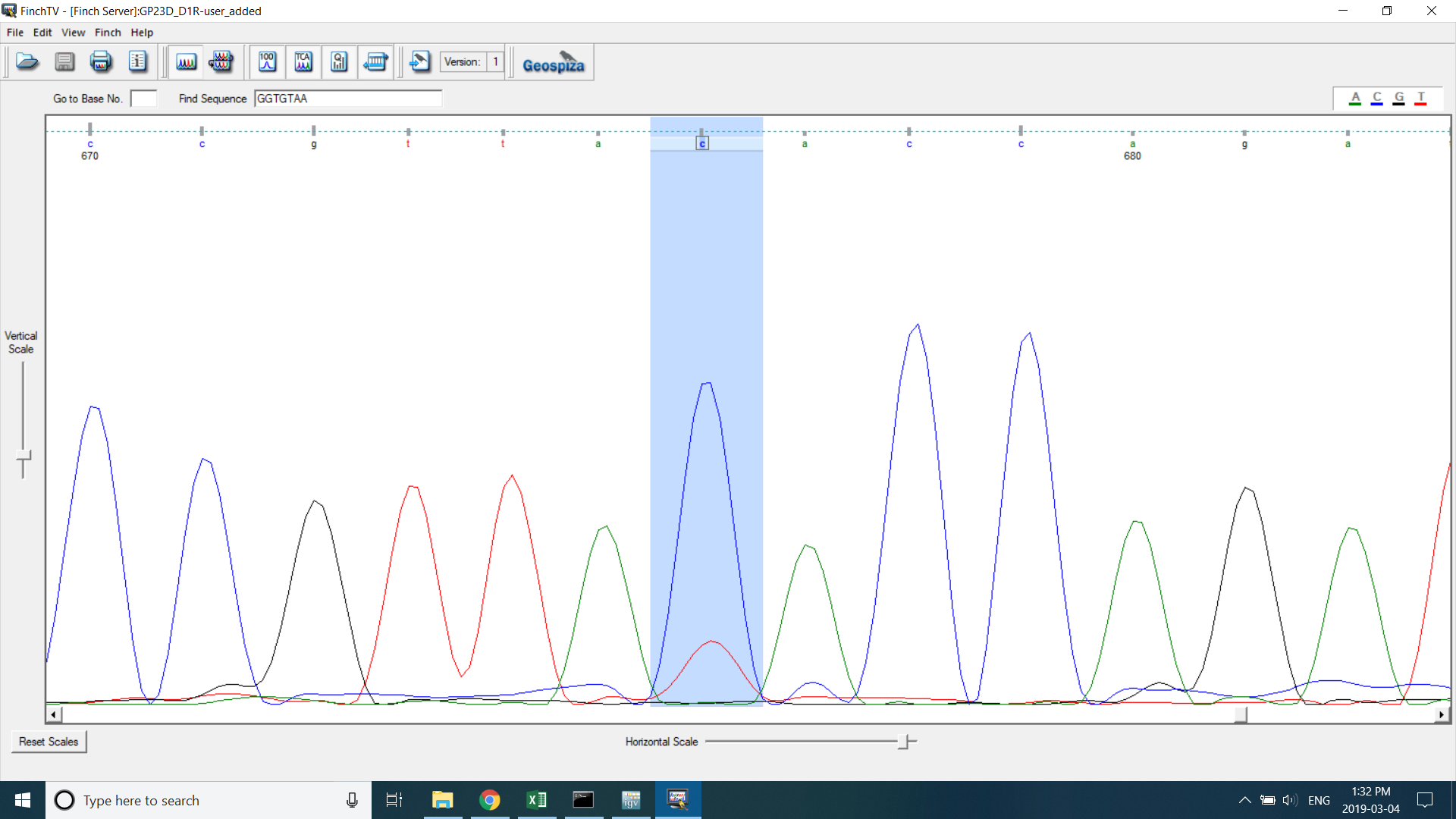

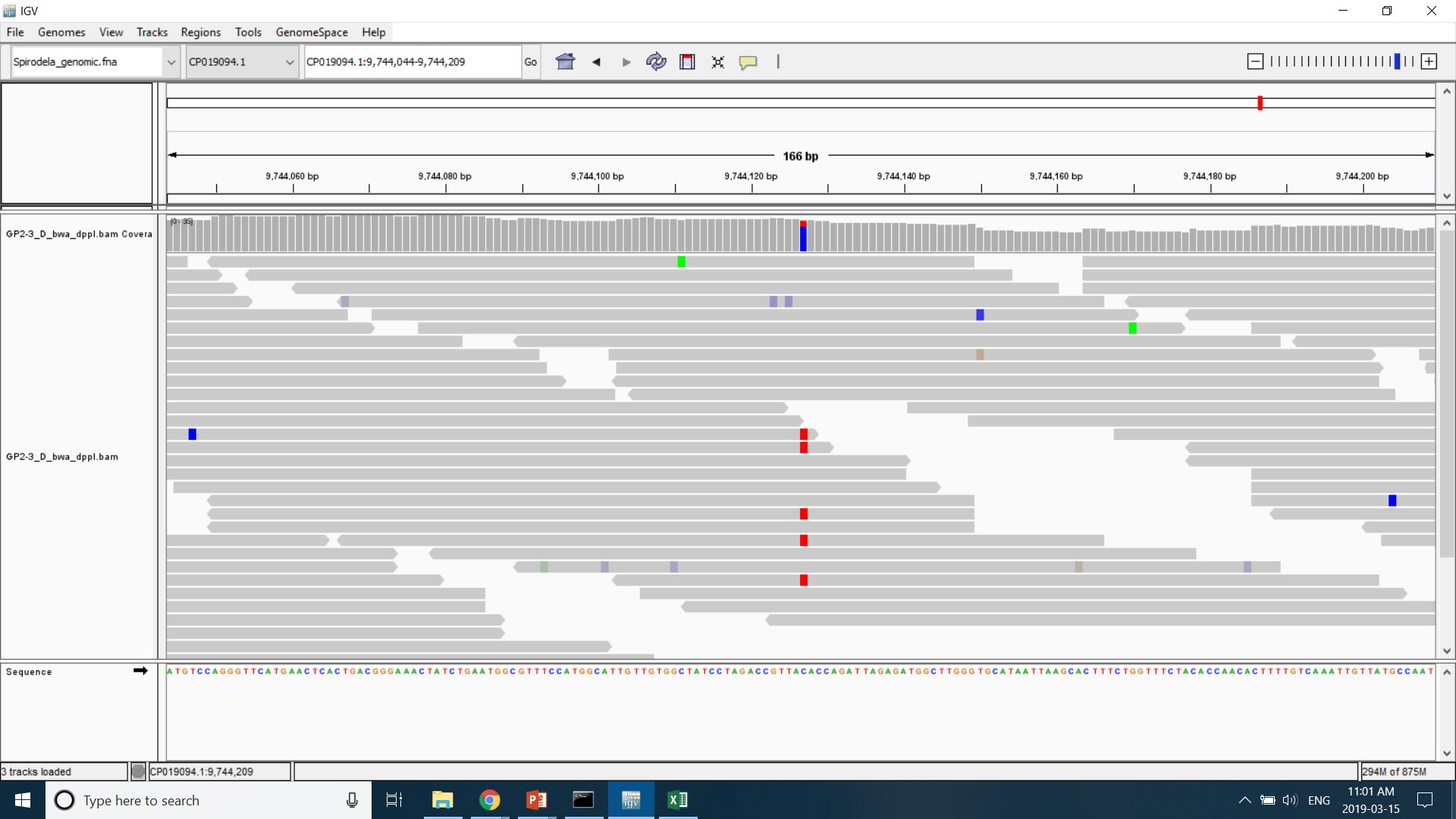

**Figure S3** Visualization of a validated *de novo* SNP from both Sanger sequencing and Illumina short read data. Both sequencing platforms suggest that the *de novo* base is present at below 50% frequency in the MA line.

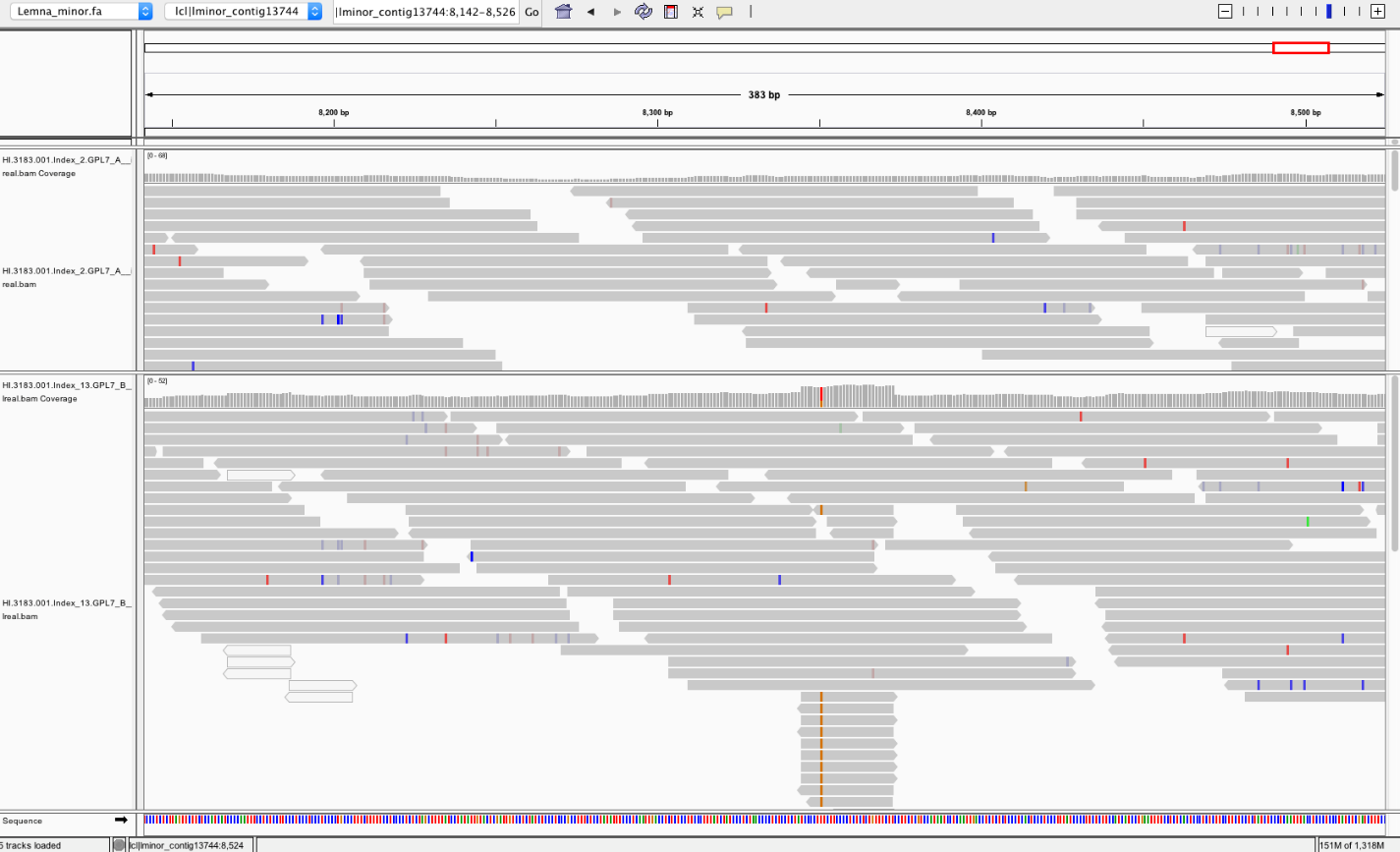
**Figure S4** Bottom track, line with a candidate *de novo* mutation that was eliminated from analysis as a suspected sequencing error. This was decided based on the short, identically mapping reads that the *de novo* base is exclusively present on. Top track, sample line not carrying evidence of a *de novo* mutation.

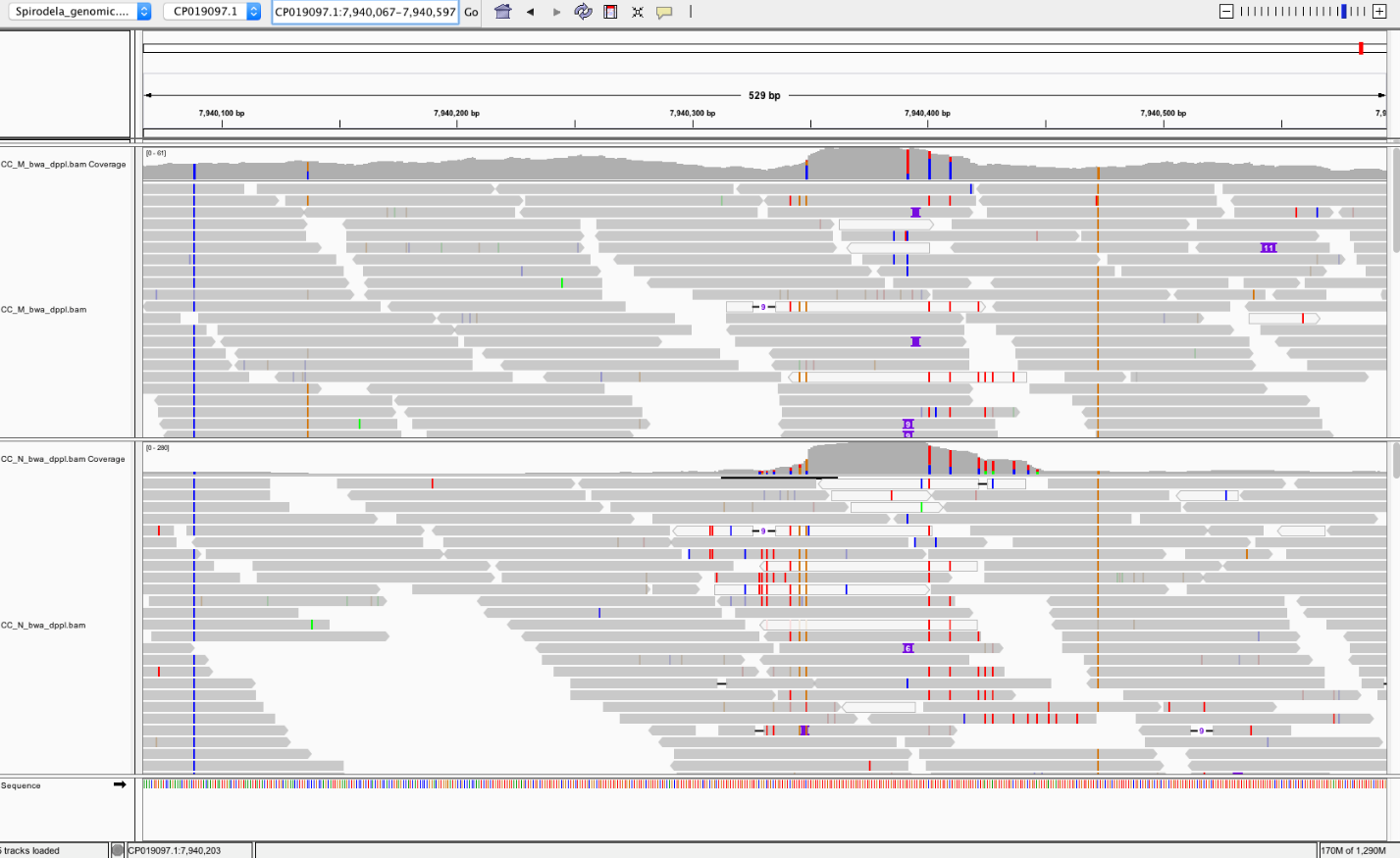

**Figure S5** Bottom track, line with a candidate *de novo* mutation that was eliminated from analysis due as a suspected error. This was decided based on the presence of highly divergent reads containing the majority of non-reference bases in this region, including the *de novo* mutant base. This is likely an issue of collapsed, divergent duplicates in the reference genome. Top track, sample line not carrying evidence of a *de novo* mutation but still showing evidence of a hidden duplication.

**Table S1** List of *de novo* mutations identified in two species of duckweed

| **Genotype** | **Scaffold** | **Position** | **Line** | **Validation** | **Reference coverage** | **Non-reference coverage** | **Site type** | **Base change** | **Effect** |
| --- | --- | --- | --- | --- | --- | --- | --- | --- | --- |
| **CC** | CP019093.1 | 4723004 | H | - | 15 | 5 | non CpG | C->G | intergenic variant |
|  | CP019097.1 | 4764451 | D | - | 14 | 5 | CpG | C->T | intergenic variant |
|  | CP019098.1 | 2797447 | P | - | 28 | 5 | CpG | C->A | missense variant |
|  | CP019100.1 | 2235140 | P | - | 19 | 5 | non CpG | C->T | intergenic variant |
|  | CP019108.1 | 782390 | N | - | 24 | 5 | non CpG | C->A | intergenic variant |
|  | CP019112.1 | 653683 | B | - | 17 | 5 | non CpG | A->T | intergenic variant |
|  | CP019095.1 | 8394653 | K | - | 26 | 6 | non CpG | C->T | intron variant |
|  | CP019105.1 | 1846849 | D | - | 15 | 6 | non CpG | A->T | intergenic variant |
|  | CP019094.1 | 7988664 | K | - | 9 | 7 | CpG | C->G | missense variant |
|  | CP019096.1 | 7777428 | K | - | 10 | 7 | non CpG | C->T | intergenic variant |
|  | CP019101.1 | 6813954 | F | - | 13 | 8 | CpG | C->T | intron variant |
|  | CP019109.1 | 2721723 | A | - | 20 | 9 | non CpG | C->T | intergenic variant |
|  | CP019093.1 | 9622614 | A | - | 18 | 11 | non CpG | C->T | intergenic variant |
|  | CP019098.1 | 7624791 | I | Pass | 9 | 13 | CpG | C->T | intergenic variant |
|  | CP019099.1 | 6962338 | C | Pass | 20 | 5 | CpG | C->T | intergenic variant |
|  | CP019101.1 | 3587320 | B | - | 25 | 5 | CpG | C->T | intergenic variant |
|  | CP019098.1 | 876453 | B | - | 15 | 6 | CpG | C->T | intergenic variant |
|  | CP019096.1 | 5852363 | D | - | 10 | 7 | CpG | C->T | intergenic variant |
|  | CP019112.1 | 199698 | P | Pass | 28 | 15 | non CpG | A->T | intergenic variant |
|  | CP019096.1 | 7993588 | C | Fail | 19 | 9 | CpG | C->T | NA |
| **GP23** | CP019094.1 | 9744127 | D | Pass | 22 | 5 | non CpG | C->T | intron variant |
|  | CP019094.1 | 10922385 | I | - | 16 | 5 | non CpG | C->T | intergenic variant |
|  | CP019095.1 | 5049789 | N | - | 35 | 5 | CpG | C->A | 5 prime UTR variant |
|  | CP019096.1 | 2739883 | P | - | 32 | 5 | CpG | C->A | intron variant |
|  | CP019103.1 | 3501136 | J | - | 13 | 5 | CpG | C->T | intergenic variant |
|  | CP019095.1 | 1536194 | N | - | 28 | 7 | non CpG | C->A | intron variant |
|  | CP019099.1 | 2234049 | D | - | 20 | 7 | CpG | C->T | intergenic variant |
|  | CP019099.1 | 5403453 | A | - | 38 | 7 | non CpG | A->G | intergenic variant |
|  | CP019103.1 | 3690446 | J | Pass | 12 | 7 | CpG | C->T | intergenic variant |
|  | CP019107.1 | 4253782 | H | - | 17 | 7 | non CpG | A->G | intergenic variant |
|  | CP019108.1 | 2476 | P | - | 30 | 7 | non CpG | C->T | intergenic variant |
|  | CP019111.1 | 3525994 | I | - | 25 | 8 | non CpG | A->G | intergenic variant |
|  | CP019108.1 | 3744394 | F | - | 10 | 9 | CpG | C->T | intergenic variant |
|  | CP019097.1 | 4835779 | P | - | 32 | 10 | CpG | C->T | intergenic variant |
|  | CP019099.1 | 6581448 | I | - | 22 | 10 | non CpG | C->T | intergenic variant |
|  | CP019104.1 | 3445982 | E | - | 19 | 10 | CpG | C->A | intergenic variant |
|  | CP019106.1 | 3255737 | O | - | 21 | 10 | CpG | C->T | missense variant |
|  | CP019093.1 | 3254558 | P | - | 44 | 12 | non CpG | C->T | splice region variant |
|  | CP019107.1 | 122485 | P | Fail | 23 | 12 | CpG | C->G | NA |
|  | CP019107.1 | 2770689 | I | - | 24 | 12 | non CpG | C->T | intergenic variant |
|  | CP019094.1 | 4157362 | A | Pass | 46 | 15 | non CpG | C->G | intergenic variant |
|  | CP019093.1 | 7920465 | H | - | 17 | 5 | non CpG | C->A | intergenic variant |
|  | CP019094.1 | 2158224 | E | - | 10 | 5 | non CpG | C->T | intergenic variant |
|  | CP019099.1 | 5294879 | J | - | 20 | 6 | CpG | C->T | intergenic variant |
|  | CP019103.1 | 2366057 | P | Fail | 46 | 5 | CpG | C->T | NA |
|  | CP019106.1 | 1753773 | A | Fail | 40 | 15 | CpG | C->G | NA |
|  | CP019106.1 | 3921984 | F | Fail | 17 | 10 | non CpG | A->G | NA |
| **GPL7** | lcl\|lminor_contig83 | 53005 | G | - | 19 | 5 | non CpG | A->C | intergenic variant |
|  | lcl\|lminor_contig645 | 31133 | A | - | 14 | 5 | CpG | C->T | intergenic variant |
|  | lcl\|lminor_contig1647 | 41425 | L | - | 14 | 5 | CpG | C->T | intergenic variant |
|  | lcl\|lminor_contig4622 | 11588 | F | - | 21 | 5 | CpG | C->T | intergenic variant |
|  | lcl\|lminor_contig7461 | 3344 | B | - | 19 | 5 | non CpG | C->T | intergenic variant |
|  | lcl\|lminor_contig7937 | 1139 | K | - | 12 | 5 | non CpG | C->T | intergenic variant |
|  | lcl\|lminor_contig12792 | 398 | O | - | 9 | 5 | non CpG | C->T | intergenic variant |
|  | lcl\|lminor_contig281 | 88447 | L | - | 17 | 6 | CpG | C->G | intergenic variant |
|  | lcl\|lminor_contig496 | 14769 | D | - | 21 | 6 | non CpG | C->T | intergenic variant |
|  | lcl\|lminor_contig560 | 25886 | A | - | 7 | 6 | non CpG | A->G | intergenic variant |
|  | lcl\|lminor_contig4164 | 4665 | A | - | 14 | 6 | CpG | C->G | intergenic variant |
|  | lcl\|lminor_contig4176 | 20576 | E | - | 10 | 6 | non CpG | C->T | intergenic variant |
|  | lcl\|lminor_contig4301 | 20542 | M | - | 16 | 6 | CpG | C->T | intergenic variant |
|  | lcl\|lminor_contig18559 | 3799 | M | - | 11 | 6 | non CpG | C->T | intergenic variant |
|  | lcl\|lminor_contig30843 | 905 | B | - | 21 | 6 | non CpG | C->T | intergenic variant |
|  | lcl\|lminor_contig1541 | 36664 | D | - | 19 | 7 | non CpG | C->T | intergenic variant |
|  | lcl\|lminor_contig4321 | 1841 | D | - | 9 | 7 | non CpG | C->T | intergenic variant |
|  | lcl\|lminor_contig4637 | 12478 | O | - | 8 | 7 | CpG | C->G | intergenic variant |
|  | lcl\|lminor_contig5967 | 746 | D | - | 15 | 7 | non CpG | C->T | intergenic variant |
|  | lcl\|lminor_contig15751 | 5324 | F | - | 9 | 7 | non CpG | C->T | intergenic variant |
|  | lcl\|lminor_contig181 | 2513 | B | - | 8 | 8 | CpG | C->T | intergenic variant |
|  | lcl\|lminor_contig5326 | 7604 | D | - | 12 | 8 | CpG | C->T | intergenic variant |
|  | lcl\|lminor_contig8410 | 1781 | K | - | 10 | 8 | CpG | C->T | intergenic variant |
|  | lcl\|lminor_contig13862 | 4502 | M | - | 15 | 8 | CpG | C->T | intergenic variant |
|  | lcl\|lminor_contig32793 | 1391 | D | - | 11 | 8 | non CpG | C->T | intergenic variant |
|  | lcl\|lminor_contig7725 | 11398 | M | - | 11 | 10 | CpG | C->T | intergenic variant |
|  | lcl\|lminor_contig2059 | 2826 | L | - | 9 | 5 | CpG | C->T | intergenic variant |
|  | lcl\|lminor_contig5191 | 11534 | D | - | 13 | 6 | CpG | C->T | intergenic variant |
|  | lcl\|lminor_contig10179 | 10642 | D | - | 18 | 10 | CpG | C->T | intergenic variant |

**Table S2** Mutation rate estimates under the observed allelic bias at *de novo* mutations

| **Predicted read freq.** | **mutation rate** | **CI lower** | **CI upper** | **Lines** |
| --- | --- | --- | --- | --- |
| 0.34 | 1.69E-10 | 8.66E-11 | 3.18E-10 | L. minor salt |
| 0.34 | 8.66E-11 | 3.47E-11 | 1.94E-10 | L. minor control |
| 0.28 | 9.91E-11 | 5.50E-11 | 1.74E-10 | S. polyrhiza salt |
| 0.28 | 8.39E-11 | 4.78E-11 | 1.44E-10 | S. polyrhiza control |

**Table S3** Mutation rate estimates under several scenarios of allelic bias at *de novo* mutant sites in salt stressed mutation accumulation lines

| **Predicted read freq.** | **mutation rate** | **CI lower** | **CI upper** | **Lines (Salt)** |
| --- | --- | --- | --- | --- |
| 0.5 | 1.26E-10 | 6.44E-11 | 2.37E-10 | L. minor (GPL7) |
|  | 9.25E-11 | 3.87E-11 | 2.01E-10 | S. polyrhiza (CC) |
|  | 6.63E-11 | 2.61E-11 | 1.50E-10 | S. polyrhiza (GP23) |
| 0.2 | 4.25E-10 | 2.17E-10 | 7.99E-10 | L. minor (GPL7) |
|  | 1.74E-10 | 7.28E-11 | 3.78E-10 | S. polyrhiza (CC) |
|  | 1.23E-10 | 4.83E-11 | 2.79E-10 | S. polyrhiza (GP23) |
| 0.1 | 3.29E-09 | 1.68E-09 | 6.18E-09 | L. minor (GPL7) |
|  | 7.47E-10 | 3.13E-10 | 1.63E-09 | S. polyrhiza (CC) |
|  | 5.00E-10 | 1.96E-10 | 1.13E-09 | S. polyrhiza (GP23) |

**Table S4** Mutation rate estimates under several scenarios of allelic bias at *de novo* mutant sites in control mutation accumulation lines

| **Predicted read freq.** | **mutation rate** | **CI lower** | **CI upper** | **Lines (Control)** |
| --- | --- | --- | --- | --- |
| 0.5 | 6.44E-11 | 2.58E-11 | 1.44E-10 | L. minor (GPL7) |
|  | 5.42E-11 | 1.99E-11 | 1.28E-10 | S. polyrhiza (CC) |
|  | 7.68E-11 | 3.61E-11 | 1.54E-10 | S. polyrhiza (GP23) |
| 0.2 | 2.17E-10 | 8.71E-11 | 4.87E-10 | L. minor (GPL7) |
|  | 1.02E-10 | 3.74E-11 | 2.40E-10 | S. polyrhiza (CC) |
|  | 1.42E-10 | 6.70E-11 | 2.85E-10 | S. polyrhiza (GP23) |
| 0.1 | 1.68E-09 | 6.74E-10 | 3.76E-09 | L. minor (GPL7) |
|  | 4.38E-10 | 1.61E-10 | 1.03E-09 | S. polyrhiza (CC) |
|  | 5.78E-10 | 2.72E-10 | 1.16E-09 | S. polyrhiza (GP23) |

**Table S5** Fraction of callable sites under various assumptions of read proportions at *de novo* mutant sites

| **Predicted read prop.** | **Callable Sites** | **Fraction Callable** | **Genotype** |
| --- | --- | --- | --- |
| No correction | 228658300 | 1.000 | GP23 |
|  | 227453248 | 1.000 | CC |
|  | 205760440 | 1.000 | GPL7 |
| 0.5 | 203374942 | 0.889 | GP23 |
|  | 203533810 | 0.895 | CC |
|  | 164847994 | 0.801 | GPL7 |
| 0.2 | 109653850 | 0.480 | GP23 |
|  | 108319605 | 0.476 | CC |
|  | 48845059 | 0.237 | GPL7 |
| 0.1 | 27002716 | 0.118 | GP23 |
|  | 25196816 | 0.111 | CC |
|  | 6317669 | 0.031 | GPL7 |
| 0.28 | 159677858 | 0.698 | GP23 |
|  | 160354540 | 0.705 | CC |
| 0.34 | 122583703 | 0.596 | GPL7 |

**Table S6** Summary of mutation rate estimates retrieved from the literature

| **Species** | **Mutation rate (per bp. per gen.)** | **Taxonomic group** |
| --- | --- | --- |
| *Chlamydomonas reinhardtii* | 3.80E-10 | Green Algae |
| *Ostreococcus tauri* | 4.79E-10 | Green Algae |
| *Ostreococcus mediterraneus* | 5.92E-10 | Green Algae |
| *Bathycoccus prasinos* | 4.39E-10 | Green Algae |
| *Micromonas pusilla* | 9.76E-10 | Green Algae |
| *Apis mellifera* | 3.4E-09 | Animals |
| *Caenorhabditis briggsae* | 1.33E-09 | Animals |
| *Caenorhabditis elegans* | 1.45E-09 | Animals |
| *Daphnia pulex* | 5.69E-09 | Animals |
| *Drosophila melanogaster* | 5.17E-09 | Animals |
| *Heliconius melpomene* | 2.90E-09 | Animals |
| *Homo sapiens* | 1.35E-08 | Animals |
| *Mus musculus* | 5.40E-09 | Animals |
| *Pan troglodytes* | 1.20E-08 | Animals |
| *Pristionchus pacificus* | 2.00E-09 | Animals |
| *Clupea harengus* | 2.00E-09 | Animals |
| *Chironomus riparius* | 2.1E-09 | Animals |
| *Ficedula albicollis* | 4.6E-09 | Animals |
| *Bombus terrestris* | 3.6E-09 | Animals |
| *Arabidopsis thaliana* | 6.95E-09 | Multicellular plants |
| *Oryza sativa* | 7.10E-09 | Multicellular plants |
| *Spirodela polyrhiza* | 8.39E-11 | Multicellular plants |
| *Silene latifolia* | 7.31E-09 | Multicellular plants |
| *Lemna minor* | 8.66E-11 | Multicellular plants |
| *Prunus persica* | 7.77E-09 | Multicellular plants |
| *Picea sitchensis* | 2.7E-08 | Multicellular plants |
| *Eucalyptus melliodora* | 6.19E-08 | Multicellular plants |
| *Neurospora crassa* | 4.10E-09 | Other unicellular eukaryotes |
| *Paramecium tetraurelia* | 1.94E-11 | Other unicellular eukaryotes |
| *Plasmodium falciparum* | 2.08E-09 | Other unicellular eukaryotes |
| *Saccharomyces cerevisiae* | 2.63E-10 | Other unicellular eukaryotes |
| *Schizosaccharomyces pombe* | 2.17E-10 | Other unicellular eukaryotes |
| *Trypanosoma brucei* | 1.38E-09 | Other unicellular eukaryotes |
| *Phaeodactylum tricornutum* | 4.77E-10 | Other unicellular eukaryotes |
